# Bioinformatic, enzymatic and structural characterization of *Trichuris suis* hexosaminidase HEX-2

**DOI:** 10.1101/2023.11.15.567194

**Authors:** Zuzanna Dutkiewicz, Annabelle Varrot, Karen J. Breese, Keith A. Stubbs, Lena Nuschy, Isabella Adduci, Katharina Paschinger, Iain B. H. Wilson

## Abstract

Hexosaminidases are key enzymes in glycoconjugate metabolism and occur in all kingdoms of life. Here, we have investigated the phylogeny of the GH20 glycosyl hydrolase family in nematodes and identified a β-hexosaminidase subclade only present in the Dorylaimia. We have expressed one of these, HEX-2 from *Trichuris suis*, a porcine parasite, and shown that it prefers an aryl β-*N*-acetylgalactosaminide *in vitro*. HEX-2 has an almost neutral pH optimum and is best inhibited by GalNAc-isofagomine. Towards N-glycan substrates, it displays a preference for the removal of GalNAc residues from LacdiNAc motifs as well as the GlcNAc attached to the α1,3-linked core mannose. Thereby, it has a broader specificity than insect fused lobes (FDL) hexosaminidases, but one narrower than distant homologues from plants. Its X-ray crystal structure, the first of any subfamily 1 GH20 hexosaminidase to be determined, is closest to *Streptococcus pneumoniae* GH20C and the active site is predicted to be compatible with accommodating both GalNAc and GlcNAc. The new structure extends our knowledge about this large enzyme family, particularly as *T. suis* HEX-2 also possesses the key glutamate residue found in human hexosaminidases of either GH20 subfamily, including HEXD whose biological function remains elusive.

## Introduction

Hexosaminidases are enzymes ubiquituous across all domains of life and play multiple roles in glycoconjugate metabolism as they remove non-reducing terminal *N*-acetylgalactosamine and *N*-acetylglucosamine residues from glycans, glycolipids, glycoproteins and glycosaminoglycans. In the case of β-hexosaminidases acting on non-reducing termini, most sequences are found in glycoside hydrolase families GH3 and GH20, which have distinct chemical mechanisms ^(*1*)^. Whereas β-hexosaminidases are primarily catabolic in mammals, in invertebrate species, they are often mediating purposeful processing steps, analogous to Golgi mannosidases, during the maturation of N-linked oligosaccharides ^(*2*)^. However, the biological significance of, and the structural basis for, hexosaminidase-mediated glycan-processing in non-vertebrates are poorly understood.

Four β-hexosaminidase genes are known from mammals: perhaps the most familiar are HEXA and HEXB encoding the α- and β-subunits of the hetero- and homo-dimeric lysosomal enzymes, which have been shown to be deficient in two storage diseases (Tay-Sachs and Sandhoff diseases, respectively) ^(*3*)^, OGA encoding a nucleocytoplasmic O-GlcNAc-specific cleaving activity of GH84 family with roles in signalling ^(*4*)^ and HEXDC encoding the nucleocytoplasmic hexosaminidase D, with an uncertain biological role. The latter enzyme ^(*5,6*)^ is a member of GH20 subfamily 1 and is only distantly related to HEXA and HEXB which are in the subfamily 2 (see ***Figure 1***). Hexosaminidase D has a neutral pH optimum and preference for aryl *N*-acetyl-D-galactosaminides ^(*5-7*)^; this enzyme is apparently significantly responsible for elevated hexosaminidase activity in synovia in rheumatoid arthritis patients ^(*8*)^.

**Figure 1:**
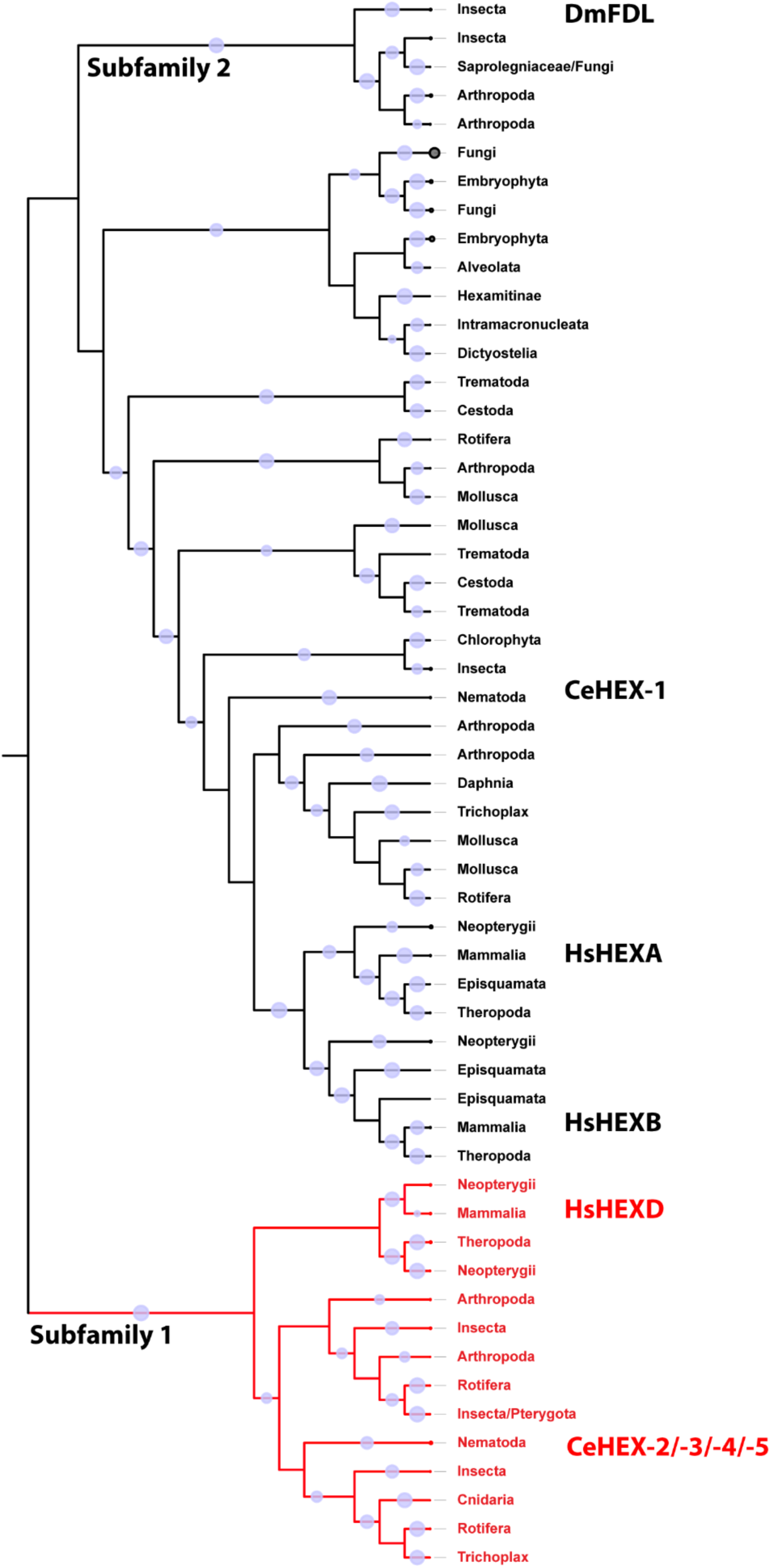
Phylogenic reconstruction of eukaryotic hexosaminidases. Subfamily 2 represents homologues of human HEXA and HEXB (HsHEXA and HsHEXB), *C. elegans* HEX-1 (CeHEX-1) and insect FDL (DmFDL); subfamily 1 (in red) contains homologues of human HEXD and *C. elegans* HEX-2/-3/-4/-5. Annotation was done based on known sequences from literature. FastTree was used to generate an approximate-maximum-likelihood phylogenetic tree, which was rooted at the midpoint. Blue circles represent bootstrap support between 70-100. Subfamilies 1 and 2 as categorized by Gutternigg et al ^(*2*)^ correspond to clades B and A defined by Intra et al ^(*28*)^.

When considering non-vertebrate species, amongst the best-described hexosaminidases are those from insects. For example, *Drosophila melanogaster* possesses a number of β-hexosaminidase genes: (i) the cytoplasmic OGA encoding the O-GlcNAc-specific enzyme similar to that in mammals ^(*9*)^, (ii) one GH20 subfamily member 1 (CG7985) with no characterised enzymatic function ^(*10*)^, but phylogenetically relatively ‘close’ to hexosaminidase D and (iii) three members of GH20 subfamily 2 including two chitinolytic and/or broad spectrum enzymes and one N-glycan-specific hexosaminidase^(*11*)^. The latter is encoded by the *fused lobes (fdl)* gene named due to the brain morphology defect in the corresponding fruitfly mutant; as enzymes, insect FDL hexosaminidases have a particular specificity for the non-reducing terminal β1,2-GlcNAc linked to the ‘lower arm’ α1,3-mannose of N-glycans ^(*11-16*)^, thereby removing the GlcNAc transferred by MGAT1 (*N-*acetylglucosaminyltransferase I). The molecular identification of FDL explained earlier work indicating that a special N-glycan-processing enzyme was present in insect cell microsomes ^(*17*)^.

The other major invertebrate model organism, the nematode *Caenorhabditis elegans*, possesses six β-hexosaminidase genes, but with a different subfamily bias as compared to insects: four of the encoded hexosaminidases belong to GH20 subfamily 1 (HEX-2, -3, -4 and -5), one to subfamily 2 (HEX-1) and one is a proven OGA from GH84 family ^(*2,18*)^. Similarly to insect FDL, HEX-2 and -3 have proven activity towards the β1,2-linked GlcNAc on the lower arm of N-glycans ^(*2*)^, corresponding to a hexosaminidase activity found in *C. elegans* microsomes ^(*19*)^; additionally, HEX-2 can also cleave non-reducing terminal GalNAc, a property also demonstrated for HEX-4 and -5 ^(*2,12*)^. HEX-1, on the other hand, is apparently chitinolytic and is close phylogenetically to human HEXA and HEXB ^(*2*)^. Our use of GFP-promoter constructs suggested different tissue expression patterns for the *C. elegans hex* genes ^(*2*)^, whereas HPLC/MS-based analyses of *hex-2*, *hex-2;hex-3* and *hex-4* mutants showed an impact of their ablation on the N-glycome ^(*2,20,21*)^. Regarding other nematodes, there is little biochemical information regarding homologous enzymes, but a secreted N-glycan-digesting hexosaminidase from *Trichinella spiralis*, with no defined sequence, has been biochemically characterised *^(22)^* and may be closest to *C. elegans* HEX-1 in terms of its properties.

Considering that *C. elegans* HEX-2 and HEX-3 are FDL-like in terms of their impact on the N-glycome, whereas HEX-4 is GalNAc-specific, we wished to explore the properties of further hexosaminidases from other nematode species. Preliminary database searching suggested that some nematodes have, like *C. elegans*, a number of GH20 subfamily 1 genes; for instance, *Oesophagostomum dentatum*, a clade V nematode like *C. elegans*, has at least HEX-2, -3 and -5 orthologues, whereas *Trichinella spiralis* and *Trichuris suis* (both clade I nematodes) appear to have only one subfamily 1 enzyme. On the other hand, *O. dentatum* lacks N-glycans with terminal GalNAc residues ^(*23*)^ and wild-type *C. elegans* has very few ^(*21*)^; both species rather have chito-oligomer-based antennae for their most complex N-glycans. In contrast, *T. suis ^(24)^* and *T. spiralis ^(25)^* are rich in N-glycans containing terminal GalNAc motifs, which may indicate a difference in the hexosaminidase-dependent processing between clade I and V species. Therefore, a thorough phylogenetic analysis was performed and the GH20 subfamily 1 candidate enzyme from *T. suis* was expressed recombinantly, characterised and successfully crystallised, yielding the first experimental structure of a eukaryotic subfamily 1 GH20 hexosaminidase.

## Results

### Phylogeny of GH20 hexosaminidases

Initially, phylogenetic analyses were performed to recreate a comprehensive evolutionary pathway for GH20 hexosaminidases, based on using almost 4,000 sequences from eukaryotes. The results verify that there are two distinct groups of these hexosaminidases: subfamily 1 and subfamily 2 (***Figure 1***). Subfamily 2 encompasses the more familiar mammalian HEXA and HEXB, as well as the insect FDL and nematode HEX-1 homologues. In subfamily 1, which includes mammalian HEXD, a clearly separated clade of nematode hexosaminidases was observed, which is represented in *C. elegans* by the previously characterized HEX-2, HEX-3, HEX-4 and HEX-5 enzymes ^(*2,12,21*)^. Each *C. elegans* hexosaminidase is within its own distinct group of related sequences from other nematodes.

In *C. elegans,* the earliest separation occurs between HEX-4 and HEX-5 (***Figure 2*** and ***Supplementary Figure 1***), indicating their specific ability to only remove GalNAc. Later, HEX-3 and HEX-2 evolved and are capable of removing both GalNAc and GlcNAc from glycan structures *in vitro* ^(*2*)^. The presence of numerous enzymes with similar functions indicates a significant amount of evolutionary pressure or possibly diverse applications for these enzymes. In general, the degree of relatedness within the GH20 clades correlates well with the proposed phylogeny of nematode species (e.g., filarial sequences are grouped together). Additionally, a subclade consisting of *Trichuris* spp. and *Trichinella* spp. representatives was identified in which only one hexosaminidase homologue per species, annotated as HEXD, was detected in the database. This finding is unusual compared to other nematodes that possess up to four subfamily 1 GH20 hexosaminidases, but reflects that *Trichuris* spp. and *Trichinella* spp. are phylogenetically distinct from the majority of nematodes, falling within nematode clade I as defined by Blaxter ^(*26,27*)^. Based on phylogenetic analysis, it can be inferred that *Trichuris* and *Trichinella* enzymes are likely to have a similar activity to that of *C. elegans* HEX-2; thus, we designated the theoretical KFD87184 sequence as *T. suis* HEX-2. Overall, *T. suis* HEX-2 has around 40% identity over ca. 500 residues with *C. elegans* HEX-2 and HEX-3 (***Supplementary Figure 2***) and shares the His/Asn-Xaa-Gly-Yaa-Asp-Glu motif with many other GH20 hexosaminidases (***Supplementary Figure 3***), whereby this sequence is shifted towards the N-terminus of subfamily 1 sequences as compared to subfamily 2.

**Figure 2:**
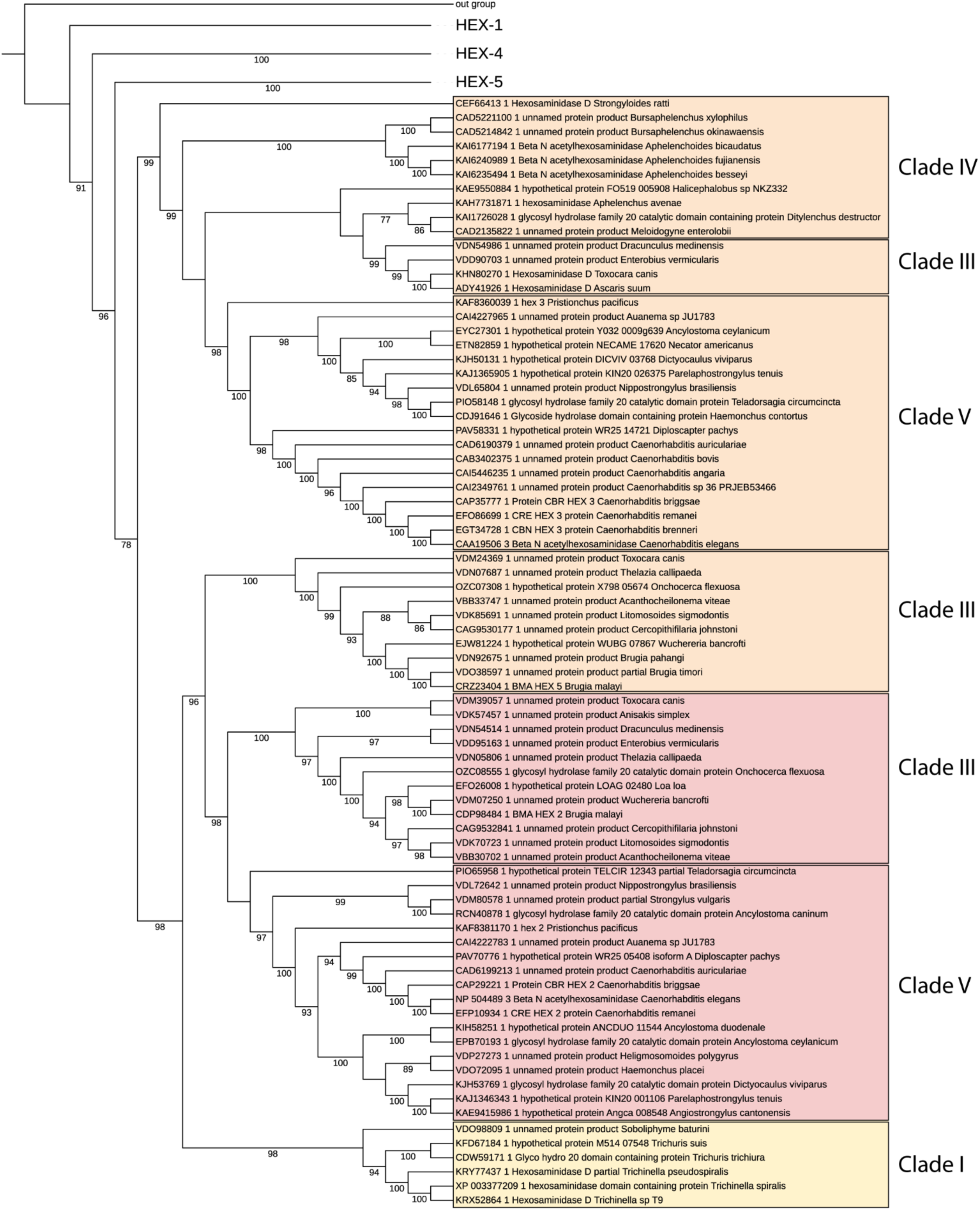
Nematode hexosaminidase phylogeny. Maximum likelihood (IQ-TREE) phylogeny of nematode GH20 hexosaminidases. The HEX-2 and HEX-3 branches are highlighted and annotated with the groups of related species in terms of the Nematoda clades as defined by Blaxter ^(*26,27*)^; ***Supplementary Figures 1A and 1B*** show all the different nematode hexosaminidase branches. The *D. melanogaster* FDL sequence was used as an out group. Bootstrap values of more than 70 are shown. In the HEX-2 branch, the sequences highlighted in yellow represent a subclade of sequences in clade I species which only have one subfamily 1 member each.

### Characterisation of HEX-2

To determine whether *T. suis* HEX-2 had an activity similar to that of *C. elegans* HEX-2, we cloned the predicted open reading frame, excluding the sequence encoding the N-terminal cytoplasmic, transmembrane and stem domains (***Supplementary Figure 2***), into a *Pichia* vector for secreted expression. Only constructs with a C-terminal His-tag could be purified by immobilized metal affinity chromatography. The resulting protein of 50 kDa (***Figure 3A***) was also verified by tryptic peptide mapping. In terms of enzymatic activity, we first examined the properties of the *T. suis* HEX-2 using artificial aryl glycoside substrates. While pNP-β-GlcNAc was a poor substrate, there was excellent activity towards pNP-β-GalNAc (***Supplementary Figure 4***). It was observed that within the linear range of product formation with respect to time and in the presence of McIlvaine buffers, optimal activity was at pH 6-7 (***Figure 3B***), similar to the values for the *C. elegans* homologues HEX-2 and HEX-4 ^(*28*)^. The optimal temperature was 60°C (***Figure 3C).*** Incubation with a range of pNP-β-GalNAc substrate concentrations and 5 ng of *T. suis* HEX-2 allowed for the determination of a *K*_m_ value of 0.89 mM (***Figure 3D***), which is also in the range for other characterized nematode hexosaminidases.

**Figure 3:**
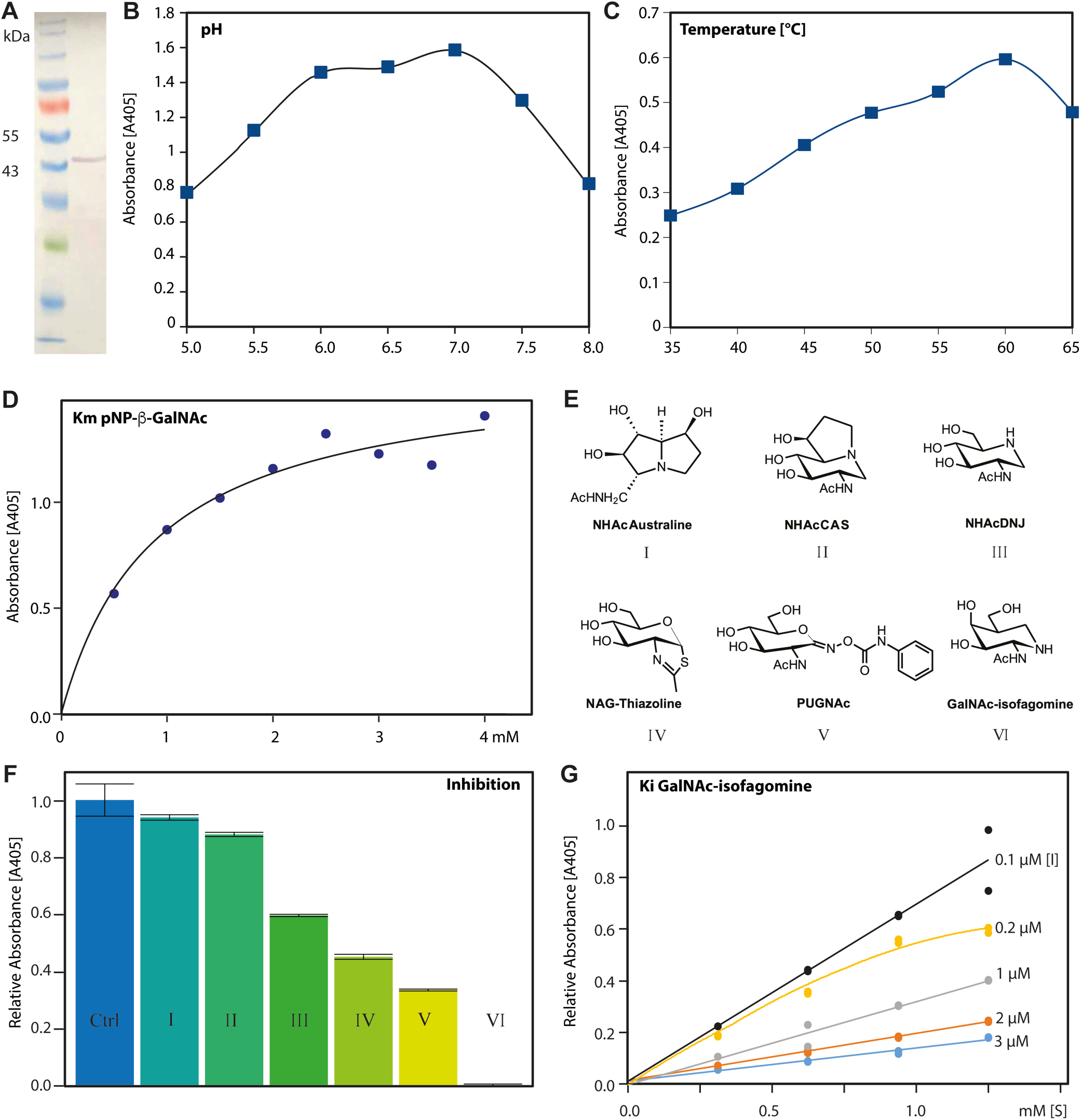
Activity of recombinant *T. suis* HEX-2 with a simple substrate. (A) Western blot of the purified recombinant ‘short’ form of *T. suis* HEX-2 expressed in *Pichia*. (B) pH dependency of activity towards pNP-β-GalNAc of recombinant *T. suis* HEX-2 assayed at 37 °C for one hour using a range of McIlvaine buffers. (C) Temperature dependency of recombinant *T. suis* HEX-2. (D) Michaelis-Menten curve for *T. suis* HEX-2 with pNP-β-GalNAc as substrate. (E and F) Competitive inhibition of *T. suis* HEX-2 protein using pNP-GalNAc as substrate (5 mM) and six different inhibitors (0.5 mM). (G) Graphical representation of the data obtained with GalNAc-isofagomine to calculate *K*_i_. Each assay was performed in duplicate or triplicate; relative absorbance is in comparison to the activity of the uninhibited enzyme.

Six different known hexosaminidase inhibitors were tested (PUGNAc, NHAcDNJ, NHAcCAS, NHAc-Australine, NAG-Thiazoline, GalNAc-isofagomine ^(*29-34*)^, ***Figure 3E***) with *T. suis* HEX-2. After pre-incubation of the inhibitors with the enzyme, pNP-β-GalNAc was again used as substrate and the activity determined. The highest degree of inhibition as compared to the control was observed with GalNAc-isofagomine and the least with NHAc-Australine (***Figure 3F***). The *K*_i_ for GalNAc-isofagomine was determined to be 0.56 µM (***Figure 3G***), a value lower than that determined for *Streptomyces plicatus* β-N-acetylhexosaminidase ^(*34*)^.

### Specificity of T. suis HEX-2

*T. suis* HEX*-*2 was tested with typical N-glycan substrates and shown to remove only one GlcNAc from a GnGn-dabsyl-N-glycopeptide (*m/z* 2060), but both GalNAc residues as well as just one terminal GlcNAc from a βGNβGN-dabsyl-N-glycopeptide (*m/z* 2467, with two LacdiNAc units) (***Figure 4 A-D***). In order to test the arm specificity, RP-HPLC analysis of a pyridylamino-N-glycan (GnGn) before and after incubation with *T. suis* HEX-2 was performed; the shift to later retention time was indicative of removing solely the ‘lower’ arm GlcNAc (***Figure 4E***) and dependent on the pH of the reaction mixture (***Supplementary Figure 3***). Another test of the specificity was to take RP-HPLC-purified core fucosylated N-glycans from *Dirofilaria immitis* with either a lower or an upper arm GlcNAc ^(*35*)^; in the case of the former, the non-reducing terminal GlcNAc was removed (***Figure 5A and B***). In contrast, *T. suis* HEX-2 did not remove the upper arm GlcNAc (***Figure 5C and D***). Regarding more complicated structures with LacdiNAc-based antennae, the activity of *T. suis* HEX-2 towards two selected N-glycan fractions derived from *T. suis* itself were selected. While the glycan with a fucosylated LacdiNAc was resistant to HEX-2 (***Figure 5E and F***), the structure with a phosphorylcholine-substituted LacdiNAc did lose the terminal GalNAc, as also reflected by the MS/MS data (***Figure 5G and H***).

**Figure 4:**
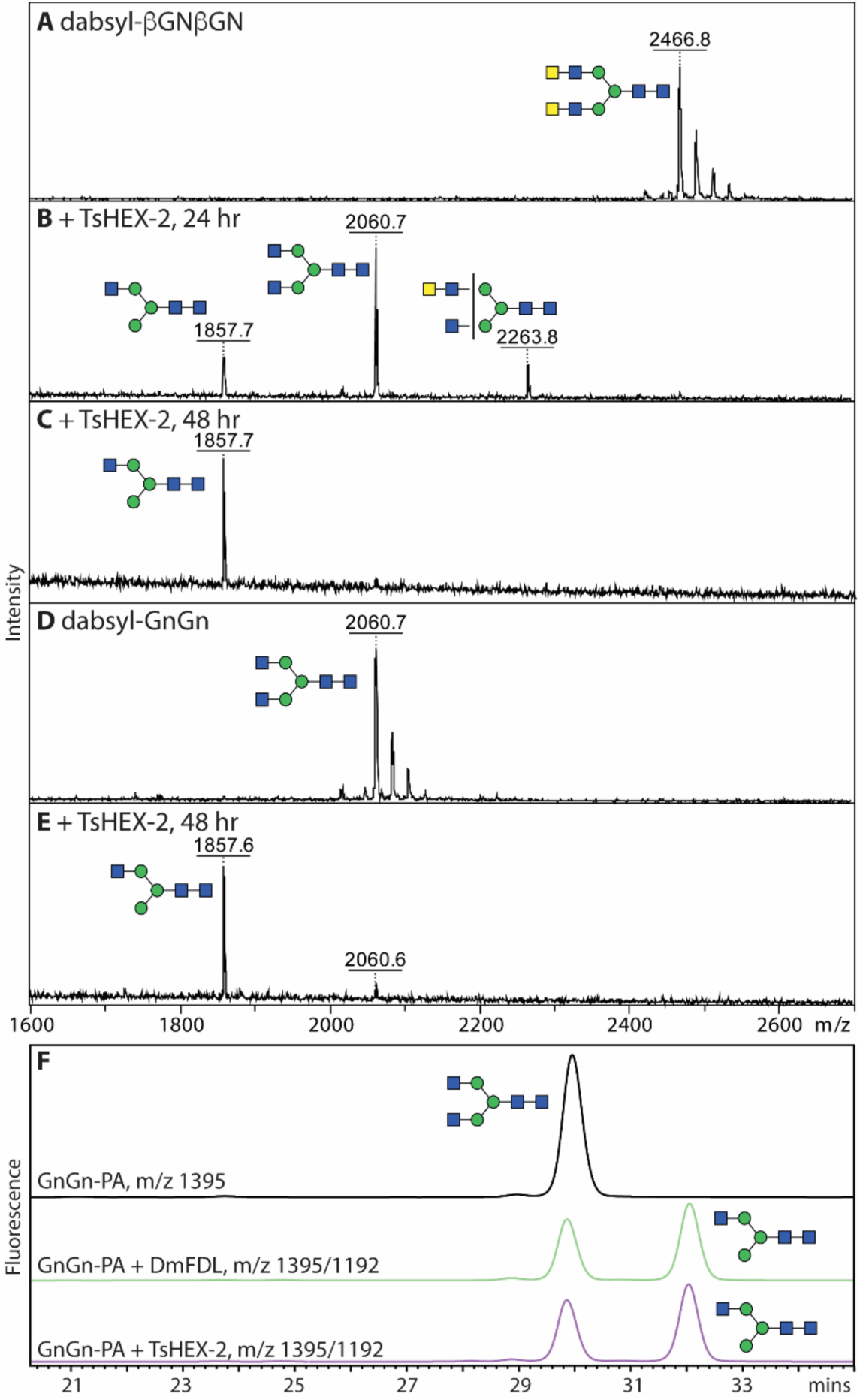
Activity of recombinant *T. suis* HEX-2 with biantennary glycan substrates. (A-E) MALDI-TOF MS analysis of incubations of dabsyl glycopeptides carrying either GnGn (*m/z* 2060) and βGNβGN (*m/z* 2467, with two LacdiNAc units) before (A/C) or after treatment with *T. suis* HEX-2 for 24 h (B), 48h (C/E). (F) RP-HPLC chromatogram of GnGn-PA (*m/z* 1395) before (black) or after treatment with either insect FDL (green) or *T. suis* HEX-2 (blue); a shift to later elution time is indicative of removal of the ‘lower’ non-reducing terminal GlcNAc residue ^(*62*)^. Red lines with arrows indicate losses of HexNAc residues from the substrates.

**Figure 5:**
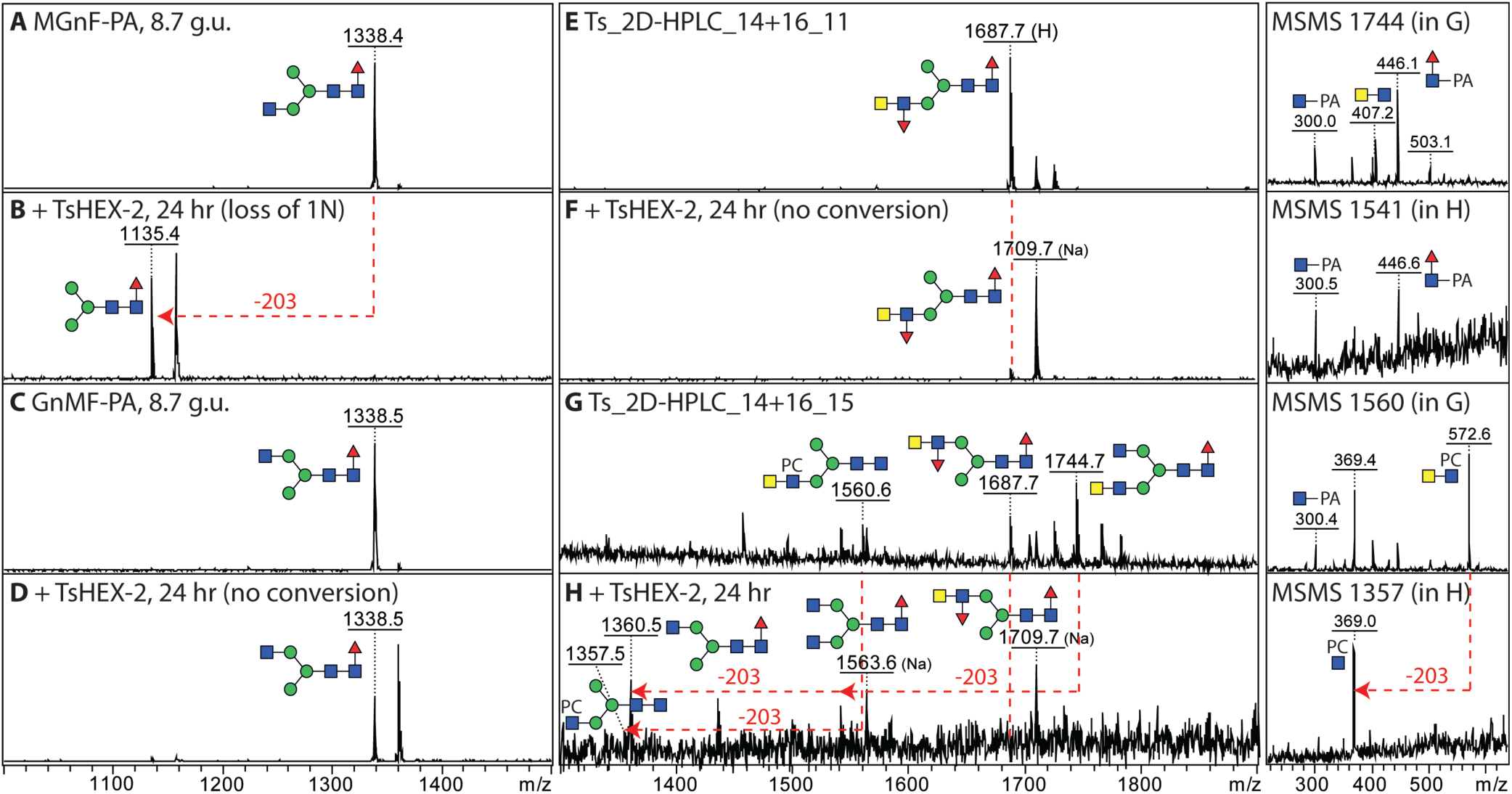
Activity of recombinant *T. suis* HEX-2 with complex nematode glycan substrates. (A-D) MALDI-TOF MS of *Dirofilaria immitis* glycans before (A/C) or after incubation with *T. suis* HEX-2 (B/D); while one Hex_3_HexNAc_3_Fuc isomer (MGnF, *m/z* 1338, eluting at 8.7 g.u. ^(*35*)^) was sensitive (B), the second isomer (GnMF, eluting at 12 g.u.) was not. (E-H) MALDI-TOF MS of *T. suis* glycans before (E/G) and after incubation (F/H) with *T. suis* HEX-2; while two glycans with fucosylated LacdiNAc motifs (*m/z* 1687 as [M+H]^+^ or 1709 as [M+Na]^+^) were not digested, structures with either non-substituted or phosphorylcholine-substituted LacdiNAc (*m/z* 1744 and 1560 as [M+H]^+^) were sensitive to *T. suis* HEX-2 (respective products of *m/z* 1563 and 1360 as [M+Na]^+^ and 1357 as [M+H]^+^), resulting in alterations in the MS/MS spectra (loss of B-ion HexNAc_2_PC_0-1_ fragments of *m/z* 407 and 572). Note that addition of HEX-2 results in a shift to sodiated ions for neutral glycans.

### X-ray Crystallography

In order to gain insight into the specificity of *T. suis* HEX-2, we solved its X-ray crystal structure at a resolution of 2.5 Å (***Table 1***). Overall, HEX-2 displays a modular structure with a N-terminal catalytic domain taking the shape of an (β/α)_8_ or TIM barrel followed by a C-terminal three helix bundle, whose function remains unknown. There is one protomer in the asymmetric unit, but a weak dimer is created by crystallography symmetry whose formation has not been confirmed in solution. Although the enzyme is predicted to be N-glycosylated, no electron density for even a core GlcNAc could be detected. Disordered regions at the N- and C- terminus correspond to the probable stem and the hexahistidine tag respectively. The structure also revealed the presence of a Zn^2+^ ion (***Figure 6A***) near the binding pocket. There is also a disordered surface loop between the Zn site and the binding side which resulted in the lack of electron density between residues Glu199 and Arg220 so they could not be modelled in the final coordinates (***Supplementary Figure 5***). Analysis of the closest related structure, the GH20C β-hexosaminidase from *Streptococcus pneumoniae ^(36)^*, indicates that this region should contain an α-helix. Additionally, the positioning of the Zn^2+^ ion near the binding pocket area implies that it could play a role in binding the substrate to the protein or in the stabilization of this disordered loop upon substrate binding. The N-terminal region of the crystallised protein with no observed electron density (residues 85-137 of the full theoretical KFD87184 sequence, corresponding to residues 1-53 of the construct) does not align well with *C. elegans* HEX-2 (***Supplementary Figure 2***) and was susceptible to proteolysis; as the ‘short’ form of the enzyme lacking this region was active, it is assumed that the stem domain extends as far as Phe138 of the full theoretical sequence.

**Figure 6:**
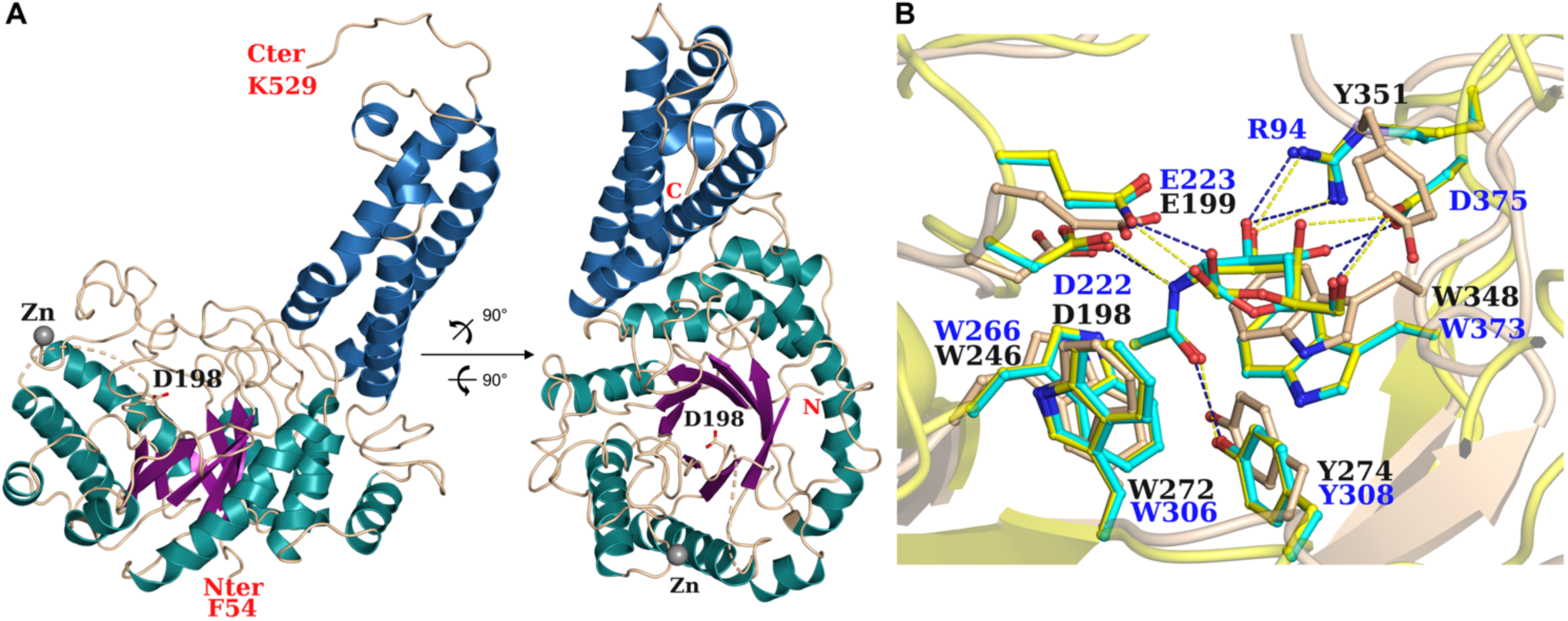
3D-structural analysis of recombinant *T. suis* HEX-2. (A) Cartoon representation of the X-ray crystal structure of *T. suis* HEX-2 at 2.5 Å in two orientations. The TIM barrel and helix bundle are colored differently, N- and C-termini with the key catalytic Asp residue represented as ball and sticks and Zn^2+^ ion as a grey ball. (B) Zoom in the binding site pocket. Overlay of *T. suis* HEX-2 structure with the structure of *S. pneumoniae* GH20C β-hexosaminidase in complex with GalNAc (yellow, PDB-ID: 5AC4) and GlcNAc (cyan, PDB-ID: 5AC5) on the catalytic domain only. Amino acids are represented in ball and sticks and H-bonds for GH20C are displayed in dash lines of corresponding color. Numbering in black is that of the recombinant secreted form for HEX-2; numbering in blue is of the full-length form of GH20C.

**Table 1:** Data collection and refinement statistics.

| Data collection |  | Refinement |  |
| --- | --- | --- | --- |
| Beamline | SOLEIL Proxima-1 | Resolution (Å) | 43.74 - 2.55 |
| Wavelength | 0.97856 | No. reflections | 17 268 |
| Space group | C2 | No. free reflections | 836 |
| Unit cell dimensions | 96.12 59.04 105.98<br>90.00 114.49 90.00 | R<sub>work</sub>/R<sub>free</sub> | 0.1894/0.2594 |
| Resolution (Å) | 43.74 - 2.55 (2.66 - 2.55) | R.m.s Bond lengths (Å) | 0.0120 |
| Nb reflections | 116 794 (14 395) | Rmsd Bond angles (°) | 1.99 |
| Nb unique reflections | 17 835 (2 157) | Rmsd Chiral (Å<sup>3</sup>) | 0.0864 |
| R<sub>merge</sub> | 0.067 (0.782) | Clashscore | 141 614 |
| R<sub>meas</sub> | 0.080 (0.931) | No. atoms/Bfac (Å<sup>2</sup>) |  |
| R<sub>pim</sub> | 0.043 (0.500) | Protein | 3 688 / 44.5 |
| Mean I/σI | 14.1 (2.1) | Water | 45 / 32.3 |
| Completeness (%) | 99.9 (99.9) | Zn | 1 / 81.3 |
| Redundancy | 6.5 (6.7) | Ramachandran<br>Allowed / Favored (%) | 99.3 / 93.4 |
|  |  | Outliers | 3 |
| CC1/2 | 0.999 (0.843) | PDB Code | 8QK1 |

As the enzymatic activity experiments indicated that *T. suis* HEX-2 removes GalNAc and GlcNAc residues from glycan substrates, the newly resolved HEX-2 structure was superimposed on that of *S. pneumoniae* GH20C (***Supplementary Figure 6***), which has been co-crystallized with the GalNAc and GlcNAc reaction products, which are also surrogates for actual substrates. This analysis indicated that either monosaccharide is capable of effectively bonding with the protein (***Figure 6B***), whereby the key contacts are conserved with the 1-hydroxyl and 2-acetamido groups. Based on the presented structure, a number of residues are predicted to participate in substrate binding within the structure (***Table 2***). These include the Asp198 and Glu199 of the His/Asn-Xaa-Gly-Xaa-Asp-Glu motif shared with other GH20 hexosaminidases (i.e., residues 282 and 283 of the full theoretical sequence), whereby the Asp and Glu are the polarising and general acid/base residues as demonstrated by studies on members of this retaining enzyme family, including a photoaffinity labelling study on human hexosaminidase B ^(*37*)^. There are though differences in terms of contacts with the monosaccharide for 3-, 4-, and 6-hydroxyls. This results a different surface loop in HEX-2 as compared to that from GH20C containing Asp375; in the *T. suis* enzyme, this loop is 5 amino acids longer and has a different conformation and sequence. Therefore, the H-bonds with the 3-hydroxyl and the 4-hydroxyl of GlcNAc are lost as well as H-bonds between 6-hydroxyl with Arg94 (***Figure 6B***). In the HEX-2 structure, those contacts are replaced by hydrophobic contacts with Tyr351.

**Table 2:**
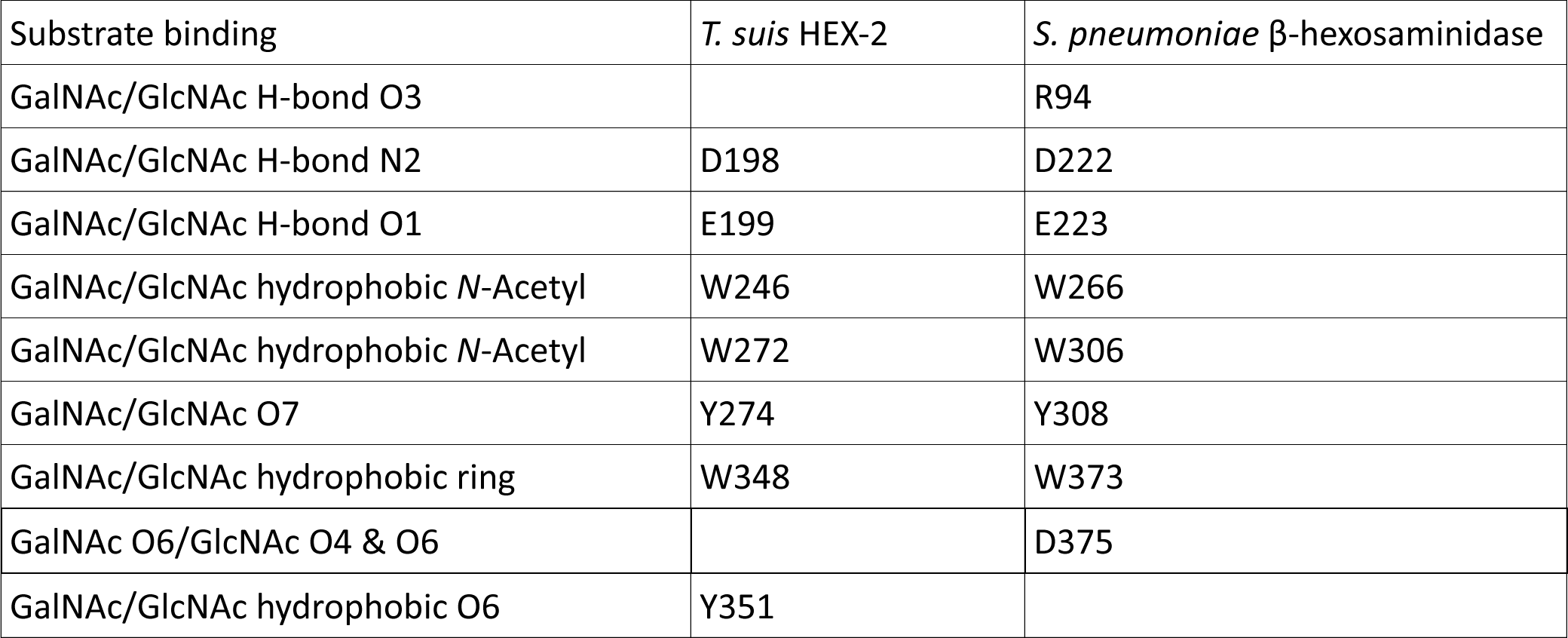
List of residues potentially involved in the binding of substrates as compared to S. pneumoniae GH20C β-hexosaminidase. Residue numbering is for the crystallised forms.

## Discussion

The phylogenetic analysis of eukaryotic GH20 hexosaminidases presented here has provided new insights into the origin of these enzymes in nematodes (***Figures 1 and 2A***). As indicated by previous studies, it was confirmed that the aforementioned HEXD is the most similar human homologue to the subfamily 1 hexosaminidases present in nematodes ^(*28*)^. Furthermore, the phylogenetic reconstruction revealed that the four major branches of subfamily 1 hexosaminidases found in nematodes derive from a single ancestor, indicating that the genes evolved through numerous duplications and later speciation. While most nematodes analysed possess multiple predicted subfamily 1 GH20 hexosaminidases, the examined clade I species (Dorylaimia; *Soboliphyme baturini*, *Trichuris* spp. and *Trichinella* spp.) have apparently only one such sequence forming a separate GH20 subbranch (***Figure 2A***).

Previously published studies have suggested that subfamily 1 GH20 enzymes prefer aryl β-*N*-acetylgalactosaminides over β-*N*-acetylglucosaminides and are generally more efficiently inhibited by galacto-epimers of hexosaminidase inhibitors as compared to those in the gluco-configuration ^(*2,7,36*)^ with our unpublished data on *C. elegans* HEX-2 also indicating a reduction in *K*_i_ for Gal-PUGNAc as opposed to PUGNAc (500-fold). In the case of *T. suis* HEX-2, no activity towards pNP-β-GlcNAc was detected, but it had a high activity with pNP-β-GalNAc and GalNAc-isofagomine was the most effective inhibitor of those tested (***Figure 3***), demonstrating its close relationship with other subfamily 1 enzymes.

The situation with N-glycan substrates is more complex than for the simple aryl glycosides: while only the ‘lower’ GlcNAc was removed from biantennary glycans with non-reducing terminal GlcNAc residues, all terminal GalNAc residues and one subterminal GlcNAc was lost from biantennary glycans with LacdiNAc motifs (***Figures 4 and 5***). This is akin to the activity of *C. elegans* HEX-2; thus, the phylogenetic designation of the *T. suis* enzyme as a HEX-2 matches its enzymatic activity. On the other hand, *C. elegans* HEX-3 only removes the ‘lower’ GlcNAc, but *C. elegans* HEX-4 is completely GalNAc-specific. Thus, despite the galacto-epimer bias for the simple substrates and inhibitors as for other subfamily 1 enzymes, *T. suis* HEX-2 can also digest a specific GlcNAc-containing linkage at slightly acidic pH in the same manner as insect FDL enzymes which are members of GH20 subfamily 2. In contrast, plant hexosaminidases which remove non-reducing terminal GlcNAc can remove both such residues from N-glycans ^(*2,12*)^.

Overall, we assume that *T. suis* HEX-2 has a role in the biosynthesis of the major paucimannosidic N-glycans known to occur in this parasite ^(*24*)^. The subtlety of its specificity towards LacdiNAc-type substrates are of interest: while *C. elegans* HEX-4 can remove GalNAc from fucosylated and phosphorylcholine-modified LacdiNAc-type motifs ^(*38*)^, the presence of an antennal fucose appears to block the action of *T. suis* HEX-2. Unlike *C. elegans*, the *T. suis* N-glycome is rich in antennal fucose motifs ^(*39*)^, but lacks the phosphorylcholine-modified chito-oligomer modifications found in a variety of other nematodes; also *C. elegans* HEX-4 appears to have a role in N-glycan biosynthesis in the Golgi apparatus and ablation of its gene leads to an increase in GalNAc-containing N-glycans in the model nematode ^(*21*)^. Thus, the fine biosynthetic control mechanism involving LacdiNAc-containing glycans may be lacking in *T. suis* and so may be a partial explanation for its species-specific glycome. On the other hand, the catabolism of GalNAc- and GlcNAc-containing N-glycans at acidic pH may be performed in *T. suis* by HEX-1, which belongs to GH20 subfamily 2; at least the potentially-related *T. spiralis* enzyme can degrade such structures ^(*22*)^.

The GH20C enzyme from *Streptococcus pneumoniae*, *^(36)^* which is the closest related protein with a crystal structure, has some bias towards GalNAc, but far less than *T. suis* HEX-2. Nevertheless, as GH20C was also co-crystallised with GalNAc, GlcNAc and some inhibitors, the superimposition with its structure is the most meaningful possible approximation; these structures were the basis for predicting potential interactions of *T. suis* HEX-2 with GalNAc and GlcNAc (***Figure 6B***). The two enzymes only superposed well on the catalytic domain (***Supplementary Figure 6***). Although the modelled conformations of the two monosaccharides are subtly different, H-bond interactions of the anomeric hydroxyl group with Glu199 (corresponding to Glu223 of GH20C) and the 2-acetamido group with Asp198 and Tyr274 (corresponding to Asp222 and Tyr308 of GH20C) would be conserved as well as the hydrophobic or stacking interactions with aromatic rings of Trp246, Trp 272 and Trp348 (corresponding to Trp residues 266, 306 and 373 of GH20C). Although the *in silico* prediction using AlphaFold was close to the model from the crystal structure (***Supplementary Figure 5***), some deviations were found, thereby highlighting the value of an experimentally-based approach. The presence of a Zn^2+^ ion near the proposed active site was not expected, its role is unknown but this is found in some other glycosidases, including Golgi mannosidase II ^(*40*)^.

Few comparisons can be made to eukaryotic GH20 hexosaminidases as the four proteins for which there is crystallographic data are all of subfamily 2. Nevertheless, the key role of an Asp-Glu pair and a Tyr residue up to 100 amino acids towards the C-terminus in binding GalNAc, GlcNAc or inhibitors is conserved ^(*41-43*)^. However, the general architecture of subfamily 2 enzymes contrasts with that of *T. suis* HEX-2, whereby the active site is closer to the N-terminus in subfamily 1. As most GH20 crystallographic studies are of bacterial enzymes, the crystal structure described here is particularly valuable as it is the first eukaryotic one from this subfamily. Thus, this structure and data presented here coupled with further studies will allow for a better understanding of the substrate specificities of invertebrate hexosaminidases and how they have evolved.

### Experimental procedures

#### Phylogenetic analyses

Eukaryotic tree: To find all eukaryotic sequences which belong to the GH20 hexosaminidase family, the Enzyme Function Initiative-Enzyme Similarity Tool (EFI-EST) was used ^(*44*)^. Two protein families IPR015883 (subfamily 1) and IPR025705 (subfamily 2) were found and all eucaryotic data was downloaded and stored on a local disk. The data has been processed to decrease the number of input sequences - any sequences below 300 or over 750 amino acids were removed. Next, sequences were used to build an alignment with MAFFT ^(*45*)^ and then the TrimAl tool was used for alignment trimming ^(*46*)^. Thereafter, the data was used as input to calculate a new phylogeny tree using the FastTree tool ^(*47*)^ on the local computer and visualized with iTOL ^(*48*)^.

Nematode tree: Characterised GH20 sequences from *C. elegans* were taken from the Wormbase database (gene names, HEX-1 – CE07499; HEX-2 – CE36785; HEX-3 – CE41720; HEX-4 – CE46668; HEX-5 – CE53609) and used as a query the whole Nematode proteome (NCBI 11.01.2023) using the hidden Markov Models algorithm from phmmer. All found sequences were used to build an alignment with MAFFT ^(*45*)^ and subsequently the final approximately-maximum-likelihood phylogenetic tree was built with IQ-tree ^(*49*)^. The resulting phylogeny tree was limited to one homologue per species and visualized with iTOL ^(*48*)^.

#### Cloning and purification

The *T. suis* HEX-2 open reading frame sequence, excluding the region encoding the cytosolic, transmembrane and stem domains (residues 1-84), was synthesized by GenScript, based on the sequence with NCBI database ID KFD67184. The hexosaminidase sequence was cloned into the pPICZαA plasmid (Invitrogen) without the native stop codon using Gibson Assembly® Cloning Kit (primers: Forward/Long: 5’- AGAGAGGCTGAAGCTGAATTCACGATGAAAGTGTATCGATGGCGA-3’, Forward/Short: 5’- AGAGAGGCTGAAGCTGAATTCACGGTGTTTATTCCGAAACGT-3’, Reverse: 5’-GACGGCACGCGTCGTATCGATAG-3’). Ligation products were transformed into NEB 5-alpha competent *Escherichia coli* (NEB #C2987) prior to selection on zeocin. The sequenced expression vectors were linearised and transformed into *P. pastoris* (GS115 strain), colonies were selected on zeocin and expression performed with methanol induction at 30 °C as previously described ^(*12*)^. The secreted long (residues 85-620) and short (residues 135-620) forms of the protein both had a C-terminal His-tag.

Protein purification was performed with an ÄKTA go protein purification system with HisTrap™ High Performance 1 ml column; samples were applied in binding buffer (25 mM sodium phosphate, 150 mM NaCl, pH 7.4) and the column washed before using a gradient of elution buffer (25 mM sodium phosphate, 150 mM NaCl, 500 mM Imidazole, pH 7.4). After purification, eluted fractions were tested for purity by SDS-PAGE; fractions of interest were pooled, concentrated using an Amicon Ultra-0.5, Ultracel-30 Membrane with 30kDa cutoff and exchanged into storage buffer (20 mM Tris-HCl, 150 mM NaCl, pH 7.5). His-tagged forms of the hexosaminidases were detectable after Western blotting using the anti-His monoclonal antibody (1:10000) and alkaline-phosphatase conjugated anti-mouse IgG (1:10000).

#### Hexosaminidase assays

The standard enzyme activity test was performed in 96 well plates. Typically, a mixture of 2.5 μl pNP-β-GalNAc (100 mM in dimethylsulphoxide), 46.5μl McIlvaine buffer pH 6.5 ^(*50*)^ and 1 μl enzyme was incubated for 1 hour at 37°C; 200 μl of stop solution (0.4 M glycine/NaOH, pH 10.4) was added and the absorbance measured with a Tecan Infinite® 200 PRO instrument. Inhibitors were prepared as previously reported ^(*30,34,51-54*)^. For tests with glycopeptides or HPLC fractions, a 1 μl aliquot was mixed with 0.2 μl enzyme and 0.8 μl of 50 mM ammonium acetate solution, pH 6.5. After overnight incubation at 37°C, 0.5 μl of the mixture was analyzed by MALDI-TOF-MS (Bruker Autoflex Speed) with 6-aza-2-thiothymine (ATT) as matrix; data were analyzed with the Flexanalysis program.

#### X-ray Crystallography

Protein crystallization was performed using commercially available crystallization screens in a hanging drop vapor diffusion setup by mixing 1 μl of protein solution (6 mg/ml) and 1 μl of crystallisation solution. The screening plate was kept in a vibration-free incubator (Molecular Dimensions, Calibre Scientific, Rotherham, UK) at 19°C. Crystal clusters were obtained from condition 18 of the Clear Strategy Screen II (Molecular dimensions) consisting of 20% PEG 1500, 0.15 M potassium thiocyanate, 0.1 M Tris pH 7.5. 15 % PEG 1000 were added to the mother liquor as cryoprotectant prior to mounting a single crystal in a cryoloop (Molecular Dimensions,) and flash freezing in liquid nitrogen. The diffraction data were collected at Synchrotron SOLEIL, Beamlines, Proxima-1 (Saint-Aubin, France) using an Eiger 16M detector (***Table 1***). The XDS ^(*55*)^ and XDSme ^(*56*)^ were used to process the data and further steps were performed with CCP4, version 8.25–27 ^(*57*)^. As crystal diffracted anisotropically, data was processed using the STARANISO server and the aimless CCP4 program. The structure of hexosaminidase was solved by molecular replacement using AlphaFold ^(*58*)^ as a search model in PHASER ^(*59*)^. Iterated maximum likelihood refinement and manual building of the resulting electron density maps were respectively performed using REFMAC 5.8 ^(*60*)^ and Coot ^(*61*)^. Five percent of the reflections were used for cross-validation analysis, and the behavior of the Rfree was employed to monitor the refinement strategy. Water molecules were added using Coot and subsequently manually inspected. The coordinates were deposited in the Protein Data Bank (PDB) under code 8QK1.

## Supporting information

PDB Validation Report

## Acknowledgements

This work was funded by the FWF-funded BioTOP doctoral programme [Austrian Science Fund W 1224] and by an FWF grant [P29466] to I.B.H.W.; I.A. is funded by the “Top Vet Science” Program of the University of Veterinary Medicine Vienna. We acknowledge the synchrotron SOLEIL (Saint Aubin, France) for access to beamlines Proxima 1 (Proposal Number 20210859) and for the technical support of Pierre Legrand. We would like to thank Valérie Chazalet and Emilie Gillon for their technical help during Z.D.’s stay at CERMAV.

## Supplemental Data

Full phylogeny of nematode hexosaminidases; predicted sequence of *T. suis* HEX-2 as well as recombinant forms and alignment of *T. suis* HEX-2 with *C. elegans* HEX-2; further data on the recombinant *T. suis* HEX-2; further views of the X-ray crystal data.

## Author contributions

Z.D. performed bioinformatic analyses and kinetics experiments; L.N., I.A., Z.D prepared constructs and purified protein, Z.D. and A.V. crystallized, collected data and solved the HEX-2 crystal structure, K.P., L.N. and Z.D. performed the glycomics experiments and interpreted data, K.J.B and K.A.S. synthesised inhibitors, Z.D. prepared draft text and figures, A.V. and I.B.H.W., funding acquisition and supervision, I.B.H.W. prepared final versions of the text and most figures. All authors have reviewed and agreed to the published version of the manuscript.

## Supplementary Information

**Supplementary Figure 1A:**
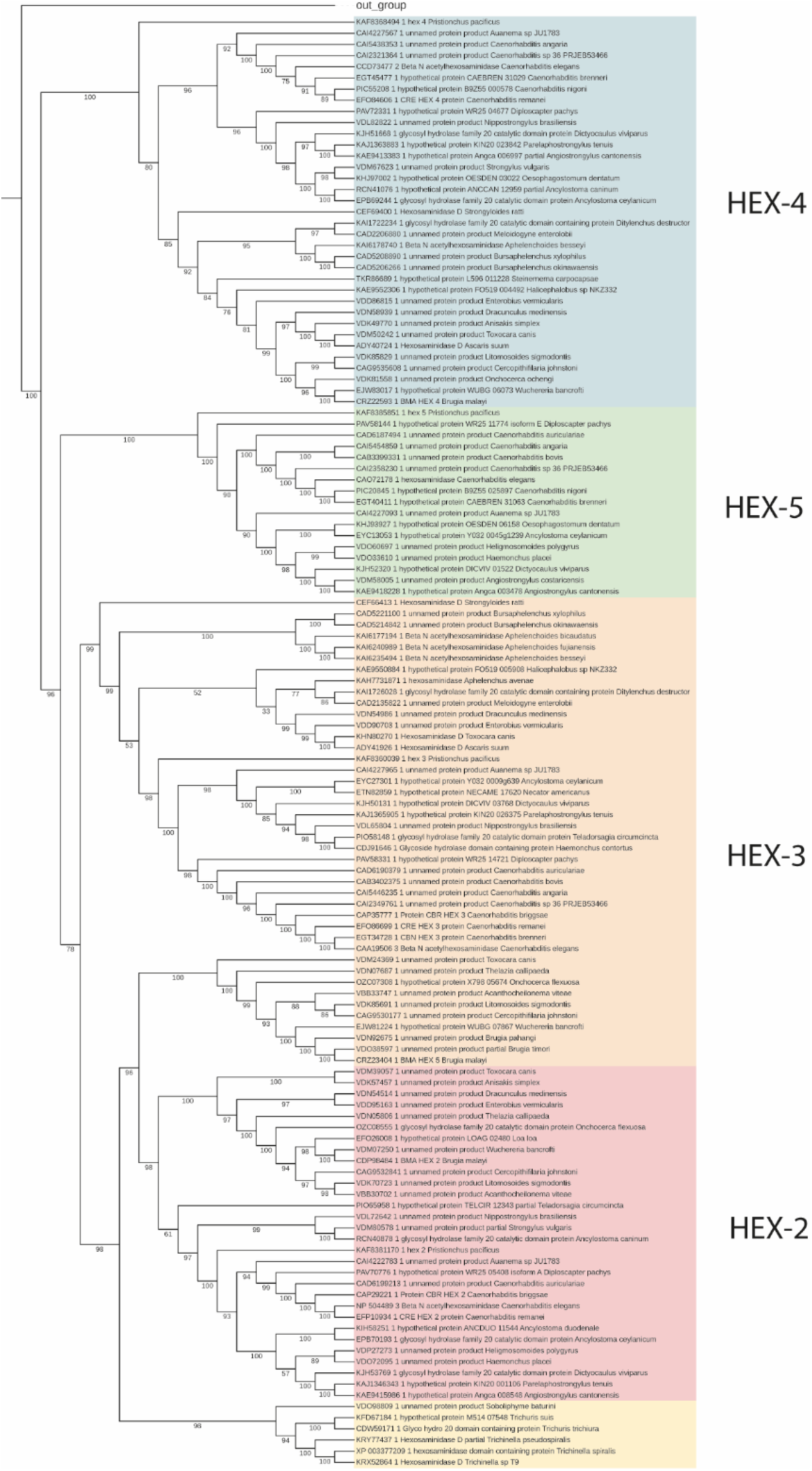
Full phylogenetic tree of nematode subfamily 1 GH20 hexosaminidases – these data underly Figure 2 in the main text. The *D. melanogaster* FDL sequence was used as an out group. Bootstrap values of more than 70 are shown.

**Supplementary Figure 1B:**
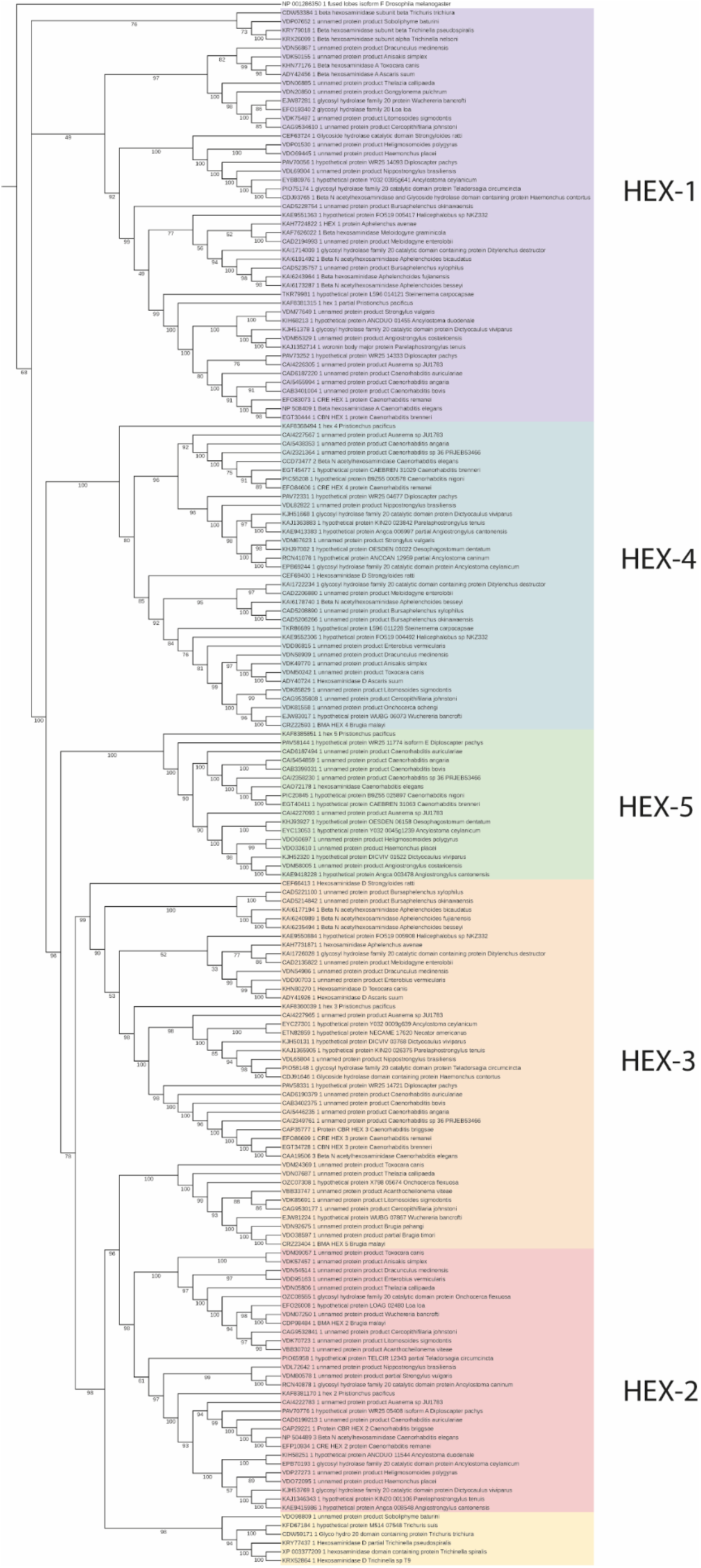
Full phylogenetic tree of all nematode GH20 hexosaminidases – these data underly Figure 2 in the main text. The D. melanogaster FDL sequence was used as an out group. Bootstrap values of more than 70 are shown.

**Supplementary Figure 2:**
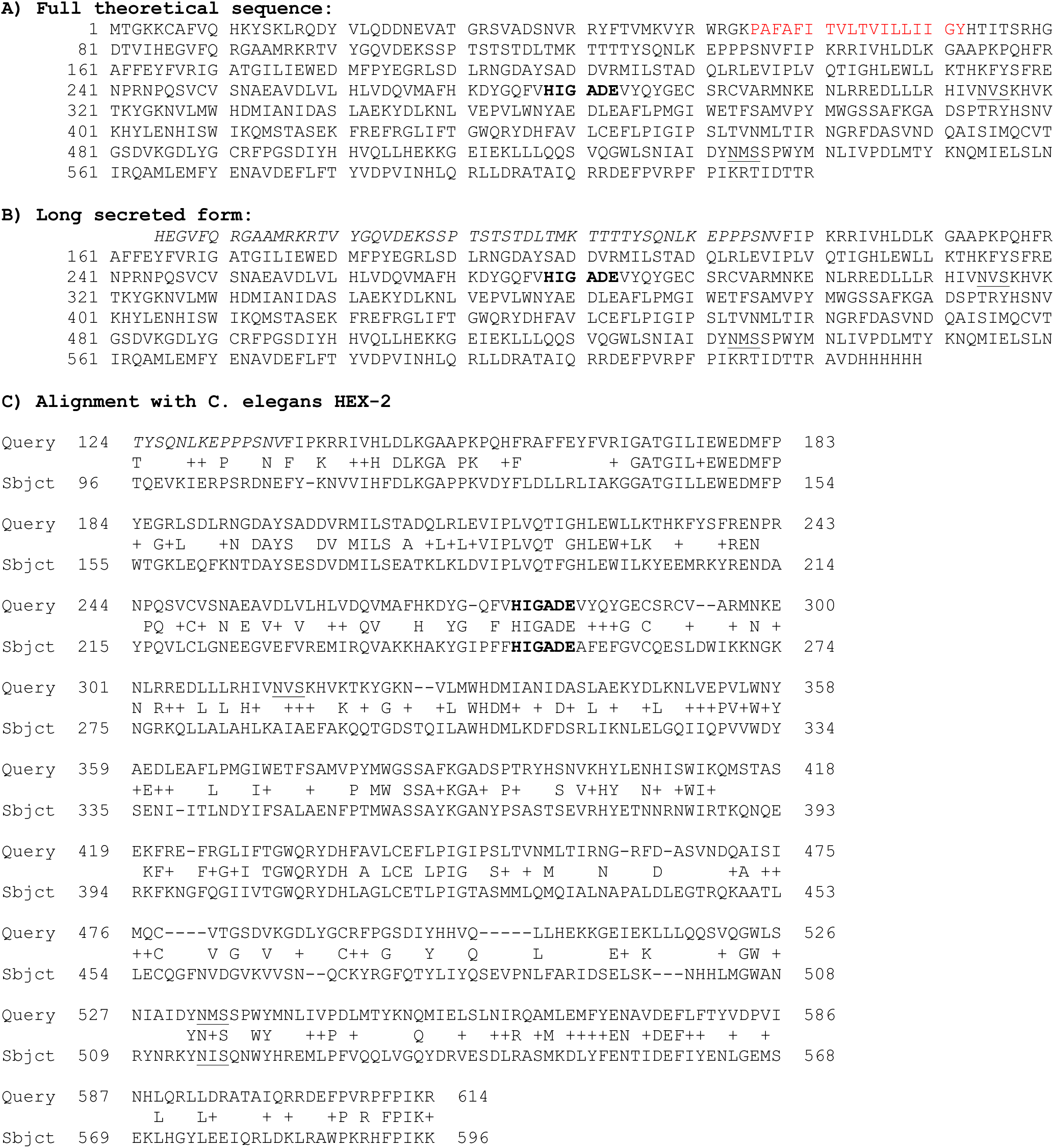
(A) Sequence of the predicted subfamily 1 hexosaminidase (HEX-2) from *T. suis.* The predicted transmembrane domain absent from the recombinant protein is in red, the potential N-glycosylation sites are underlined and the conserved HIGADE active site region is indicated in bold; the actual initial methionine residue is unknown but two are N-terminal to the transmembrane domain. (B) Sequence of the recombinant long secreted form (expected M_r_ 63 kDa) used for X-ray crystallography; the N-terminal disordered region absent from the recombinant short secreted form (expected M_r_ 58 kDa) is italicized. (C) Alignment of *T. suis* and *C. elegans* HEX-2.

**Supplementary Figure 3:**
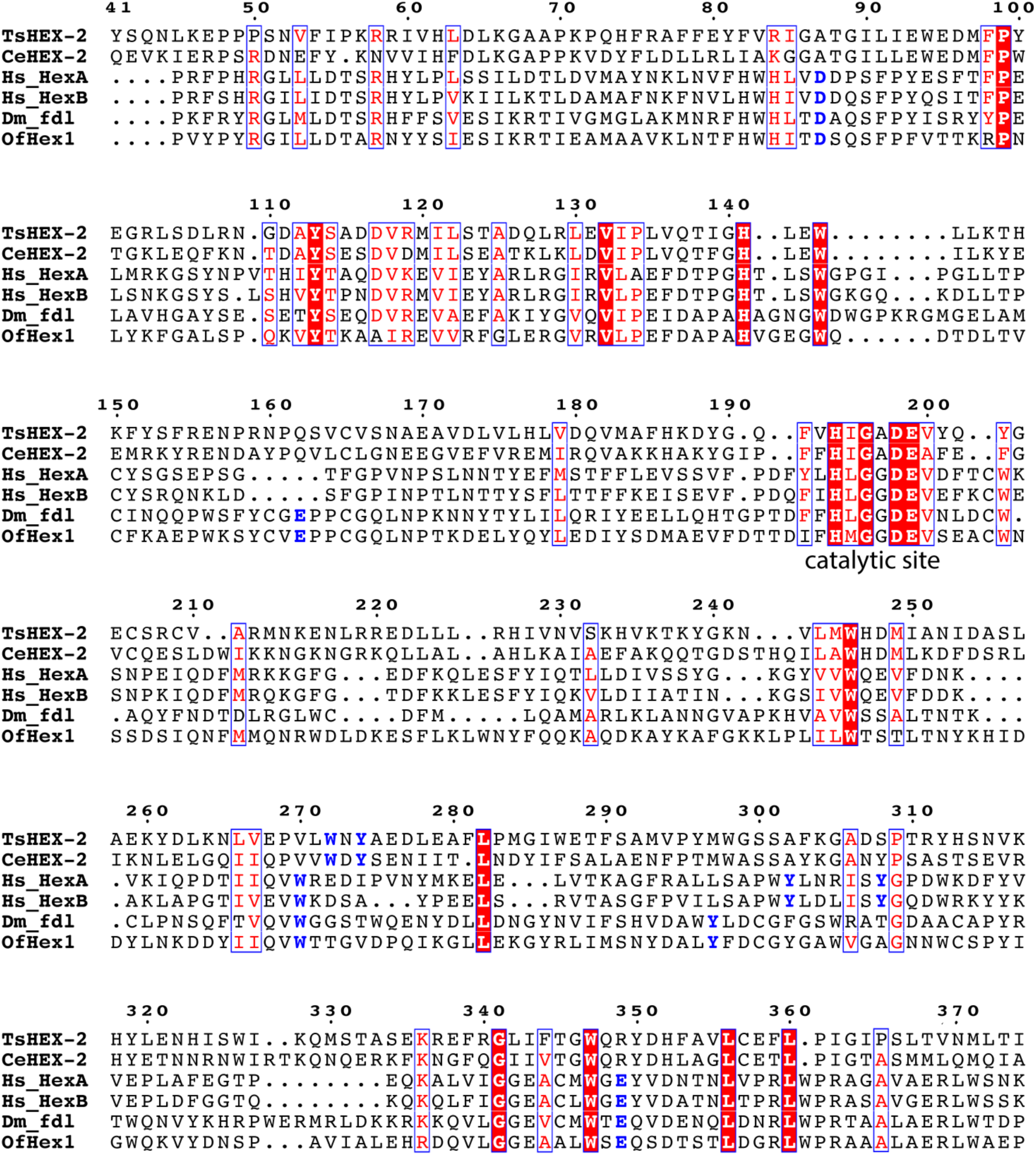
Alignment of *T. suis* and *C. elegans* HEX-2 (both subfamily 1), human HEXA and HEXB, *Drosophila melanogaster* FDL and *Ostrinia furnacalis* OfHex1 (subfamily 2). Sequences were shortened to the region of highest homology and aligned using Multalin, http://multalin.toulouse.inra.fr/ #x002D; the file was processed on https://espript.ibcp.fr/ESPript/ESPript/ Crystal structures exist for human hexosaminidases A and B as well as OfHex1, while HEX-2 and FDL specifically remove the β1,2GlcNAc from the α1,3-mannose of N-glycans. Only the catalytic site as well as a region towards the N-terminus is well conserved between the two subfamilies. Highlighted in blue are selected key subfamily-specific residues identified in this and other structural studies (other than those conserved across the families in red).

**Supplementary Figure 4:**
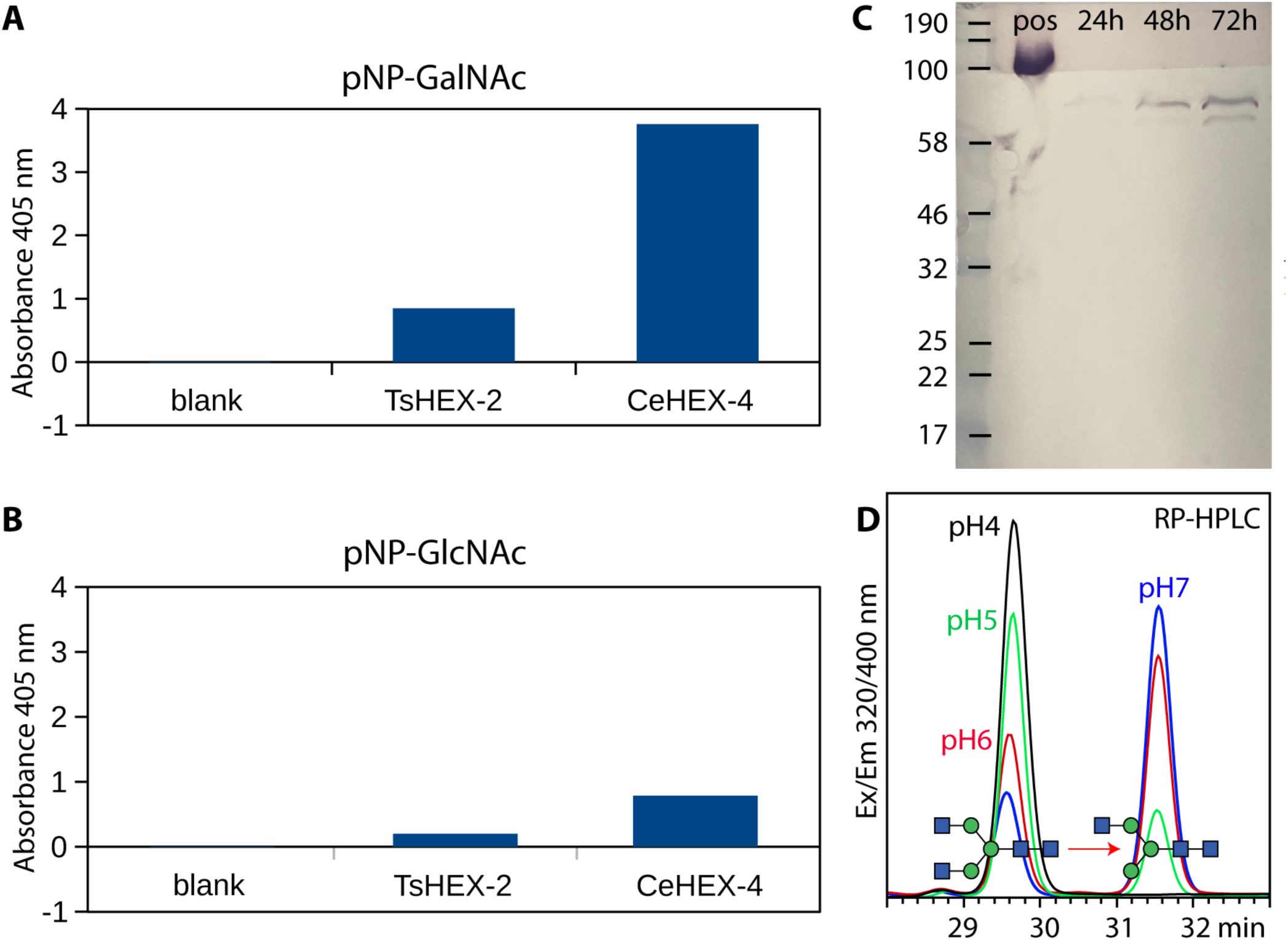
(A and B) Activity towards pNP-β-GalNAc and pNP-β-GlcNAc of *T. suis* HEX-2 and *C. elegans* HEX-4 expressed in *Pichia pastoris*; the turnover of the aryl GalNAc substrate is some three- or fourfold higher than that of the aryl GlcNAc in keeping with data on other subfamily 1 GH20 hexosaminidases. (C) Expression of the recombinant secreted ‘long’ form of *T. suis* HEX-2 (expected M_r_ 63 kDa) as detected by Western blotting of *Pichia* culture supernatants after 24, 48 or 72 hours of induction. (D) Activity of *T. suis* HEX-2 at different pH values towards GnGn-PA; the appearance of the GnM-PA product of later RP-HPLC elution time (31.5 mins; see also Figure 4 of the main text) is most pronounced at pH 6 and 7, in keeping with the optimum found with the aryl GalNAc substrate (see Figure 3 of the main text).

**Supplementary Figure 5:**
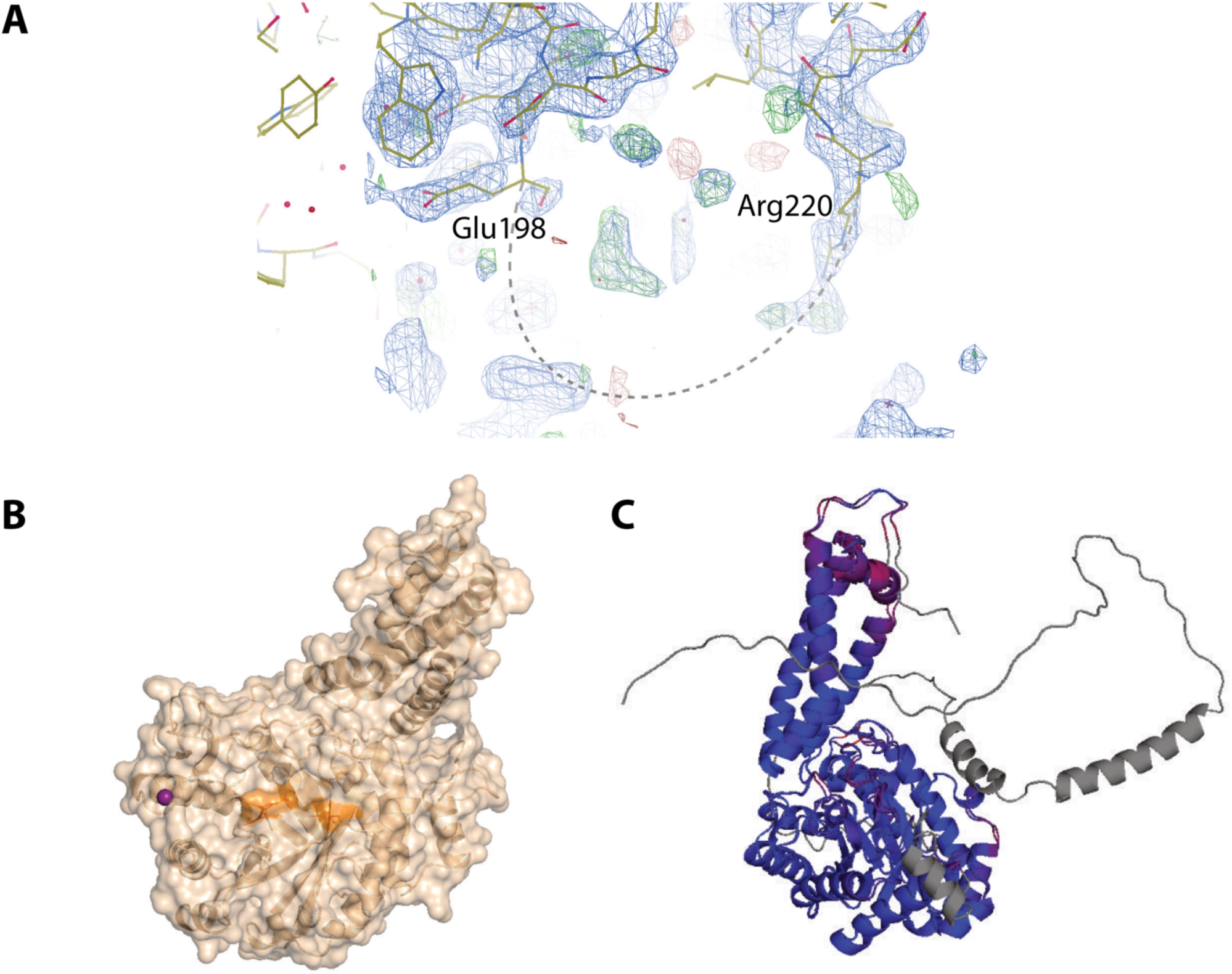
(A) Electron density maps in the disordered area of the *T. suis* HEX-2 structure, visualized with Coot (version 0.9.8.3). Difference map Fo-DFc map is represented in green (positive) and red (negative) at 3 sigmas whilst 2Fo-DFc is represented in blue at 1 sigma (0.23 eÅ^3^). (B) Cartoon and surface representation of crystal structure of *T. suis* HEX-2 at 2.5 Å; the purple ball represents the Zn^2+^ ion and 2 presumed catalytic amino acids are displayed in lines and orange colored. (C) Comparison of the *T. suis* HEX-2 X-ray structure and the Alpha Fold model, coloured by RMSD (blue specifying the minimum pairwise RMSD and red indicating the maximum, unaligned residues are coloured grey), visualized with pymol (free version 2.5.0, Schrodinger LLC). See also Figure 6 of the main text for other visualisations.

**Supplementary Figure 6:**
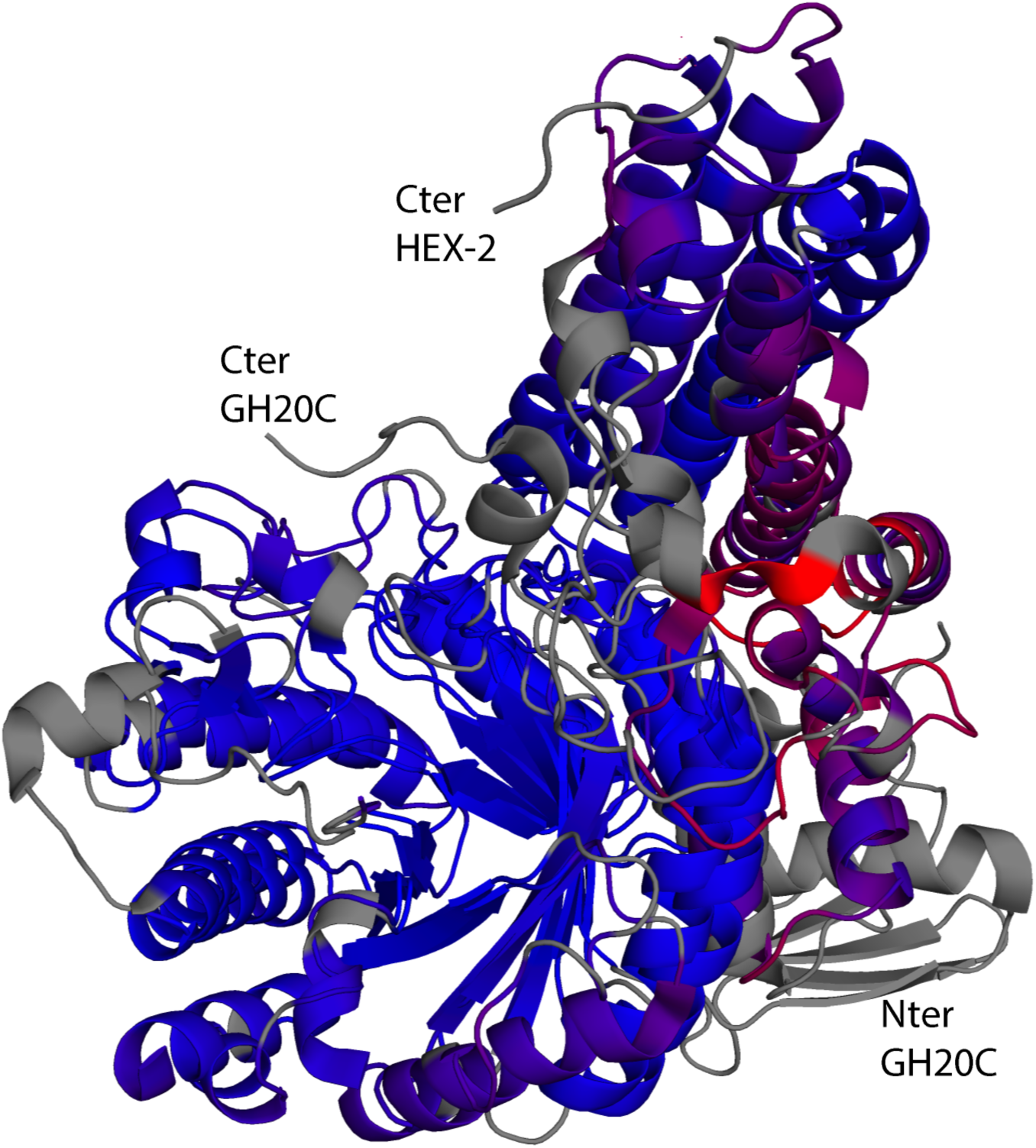
Superimposition of *T. suis* HEX-2 (8QK1) and *S. pneumoniae* GH20C (5A6J) monomeric crystal structures coloured by RMSD as visualized with pymol (free version 2.5.0, Schrodinger LLC). The distances between aligned C-alpha atom pairs are stored as B-factors of these residues, which are coloured by a spectrum ranging from blue specifying the minimum pairwise RMSD and red indicating the maximum. Unaligned residues are coloured grey. The barrels containing the binding pocket superimpose well, but the N- and C-termini differ (in this view, the N-terminus of HEX-2 is hidden under the barrel). See also Figure 6 of the main text for other visualisations.

